# Genomics of Irish swine-derived *Streptococcus suis*: Population structure, prophages, and anti-viral defence mechanisms

**DOI:** 10.1101/2025.08.29.672906

**Authors:** Emmanuel Kuffour Osei, A. Kate O’Mahony, Reuben O’Hea, John Moriarty, Áine O’Doherty, Margaret Wilson, Edgar Garcia Manzanilla, Jennifer Mahony, John G Kenny

## Abstract

*Streptococcus suis* is a major pig pathogen with zoonotic potential, posing an occupational risk to farmers and meat handlers. We characterised 110 *S. suis* strains from diseased pigs in Ireland (2005–2022) using whole-genome sequencing to investigate population structure and phage-host dynamics. We identified fifteen distinct serotypes, with serotypes 9 and 2 being the most dominant. *In silico* multi-locus sequence typing revealed high diversity within the collection, identifying several sequence types (STs), including 26 novel STs. Investigation of strain-level genomic clustering using PopPUNK against global *S. suis* genomes showed that the Irish isolates were phylogenetically dispersed across the broader global *S. suis* population rather than clustering in a single clonal group. The majority of Irish isolates fall within the ten established pathogenic lineages, including the highly virulent zoonotic lineage 1. A stable endemic clonal lineage was identified among Irish isolates, showing minimal genetic variation over a decade.

Prophage analysis revealed novel viral taxa that were interspersed among known streptococcal phages, rather than clustering distinctly. Restriction-modification systems were the predominant anti-viral defence systems identified across genomes. CRISPR-Cas systems were present in limited strains but showed substantial targeting bias toward full-length prophages, indicating ongoing phage pressure. CRISPR spacers matched non-*S. suis* streptococcal phages, and phylogenomic analysis revealed that *Vansinderenvirus* phages clustered with *S. suis* rather than other *S. thermophilus* phages, suggesting evolutionary connections between phage lineages infecting different streptococci.

This study presents the first comprehensive genomic characterisation of *S. suis* in Ireland, revealing a diverse population with significant implications for animal and human health.

## Introduction

Pathogens often emerge by crossing the species barrier; however, they can also arise from adaptive evolution within the host microbiota through horizontal gene transfer, recombination, and mutation, enabling the acquisition of virulence factors (Murray et al., 2023). One such bacterium is *S. suis*, which typically colonises the upper respiratory tract of pigs as part of the normal microbiota. Under conditions such as post-weaning stress and where co-morbidities exist, *S. suis* can cause respiratory and systemic infections such as meningitis, polyarthritis, and endocarditis (Li et al., 2024). These infections result in significant production losses with economic costs of up to €14,000 annually per production unit and mortality rates reaching 20%, alongside animal welfare concerns including severe pain from arthritis and meningitis (Neila-Ibáñez et al., 2021). Beyond infections in pigs, *S. suis* is a zoonotic pathogen and has caused three deadly outbreaks in humans since 1998 (Dong et al., 2021). Human infections mediated by *S. suis* occur through direct contact with infected pigs, posing occupational risks to farmers and animal and meat handlers throughout the farm-to-fork continuum, and via consumption of raw or undercooked pork products. For surveillance and diagnostic purposes, the bacterium is classified into 29 serotypes based on capsular polysaccharides (CPS) and over 2800 sequence types (ST) based on seven housekeeping genes (King et al., 2002; Li et al., 2024). Although there are spatiotemporal variations in the serotypes and STs, isolates of serotype 9 ST16 and serotype 2 ST1 are frequently implicated in pig diseases in Europe (Goyette-Desjardins et al., 2014). The distribution of *S. suis* genotypes in pig and zoonotic infections globally have been reviewed elsewhere (Goyette-Desjardins et al., 2014; Segura et al., 2020).

In Ireland, *S. suis* could potentially pose a similar concern for the pig industry and, though rare, for human health. While there is no comprehensive genomic surveillance data in Ireland, veterinary reports indicate that *S. suis* is the most common cause of meningitis in pigs submitted to diagnostic laboratories (Department of Agriculture, Food and the Marine, 2021; MSD Animal Health, 2022). Moreover, we recently reported that meningitis accounted for 41.8% of all isolates recovered from pig carcasses in Ireland, with confirmed *S. suis*-associated disease between 2010 and 2024 (Moriarty et al., 2025). Globally, control strategies rely heavily on antibiotics, although their long-term sustainability is called into question by the rising threat of antimicrobial resistance (AMR). Additionally, while autogenous vaccines are effective at the level of application in individual farms, their broader effectiveness is confounded by the antigenic diversity of *S. suis* (Segura et al., 2020). Candidates for off the shelf vaccines, such as CPS, offer limited cross-protection, particularly when multiple serotypes or divergent strains co-circulate within a herd. An additional hurdle to the control of *S. suis* is the fact that it is not a notifiable disease in several countries, which limits surveillance data and hinders the development of coordinated, sector-wide responses (Osei et al., 2022; Segura et al., 2020). Misidentification and misdiagnosis are not uncommon due to the use of traditional serological typing methods such as slide agglutination tests that cannot resolve the genetic subtleties that distinguish pathogenic streptococci species or strains (Chen et al., 2016; Meekhanon et al., 2019; Tarini et al., 2019).

Together, these challenges necessitate the implementation of genomic epidemiological surveillance to monitor the dissemination and evolution of *S. suis* and to guide the development of more effective, broadly applicable interventions to enhance food safety. In this study, we present the first population-level genomic investigation of *S. suis* in Ireland, based on isolates recovered from diagnostic pig carcass submissions from 2005 to 2022. Using whole-genome sequencing, we characterise the population structure of Irish isolates and contextualise them in the global *S. suis* landscape. In parallel, we investigated the interplay between prophages and anti-viral defence systems in relation the evolutionary dynamics of the *S. suis* population. The interactions between prophages and defence systems highlight an active phage-host interface that contributes to genome plasticity and may shape ecological success and community dynamics of *S. suis* within the pig microbiota. Exploring these dynamics is key to understanding pathogen evolution and carries a broader significance from a One Health perspective, given the growing interest in phage-based controls for foodborne and zoonotic pathogens.

## Materials and Methods

### Bacterial isolation, identification and cultivation

Between 2005 and 2022, swab and post-mortem tissue samples collected from pig carcasses diagnosed with *S. suis* infections were processed and plated on sheep blood agar at the Veterinary Laboratory Service of the Department of Agriculture, Food and the Marine Laboratory, Backweston, Ireland (Moriarty et al., 2025). Preliminary identification of *S. suis* was carried out using matrix-assisted laser desorption ionisation time-of-flight mass spectrometry (MALDI-TOF MS) with a cut-off score of 2.0, according to manufacturer’s instruction (Bruker Daltronik GmbH, Bremen, Germany). In total, 112 presumptive *S. suis* isolates were received at Teagasc Food Research Centre for further characterisation.

### Molecular serotyping and whole-genome sequencing

A two-step multiplex PCR was performed to classify *S. suis* isolates into serotypes as previously described (Okura et al., 2014). Genomic DNA was extracted from pure overnight cultures grown in Todd Hewitt broth using the Monarch Genomic DNA Purification Kit (New England Biolabs, #T3010L) according to the manufacturer’s instructions for gram-positive bacteria DNA purification. The first round of PCR (grouping PCR) classified the isolates into one of seven groups based on their capsular polysaccharide genotype (*cps*), and the second step determined the specific serotype within each group. For whole-genome sequencing, genomic DNA from *S. suis* isolates was quantified using the Qubit 1X dsDNA HS kit (Invitrogen, #Q33231). DNA was then normalised and sheared to ∼350 bp fragments by sonication, followed by end-repair, dA-tailing, and ligation of Illumina-compatible adapters. Library quality was assessed using Qubit 2.0 and Agilent Technologies 2100 Bioanalyzer. Uniquely indexed libraries were pooled and sequenced on a NovaSeq 6000 (S4 flow cell) by Novogene Ltd, generating ∼1 Gb of 2 × 150 nt paired-end data per sample. The raw sequence reads have been deposited in the NCBI Sequence Read Archive and are available under BioProject PRJNA1295417

### Genome assembly and annotation

Trim galore (v0.6.1), which uses FastQC (v0.11.8) and Cutadapt (v2.6), was used for checking raw reads quality, trim low-quality base calls and remove adapter sequences from 3’ ends of reads (https://github.com/FelixKrueger/TrimGalore). *De novo* assemblies were generated using SPAdes (v3.15.3) and the quality metrices of assembled genomes including N50, contigs per genome, GC content, and genome size were examined using QUAST (v5.1.0) (Gurevich et al., 2013; Prjibelski et al., 2020) (Table S1). To assess genome completeness and contamination, the “lineage_wf” workflow in CheckM (v1.0.18) was used (Parks et al., 2015). Taxonomic assignment was confirmed using the gtdb_tk (v2.4.0) toolkit (Chaumeil et al., 2020) with database release220. Prokka (v1.14.5) was to used annotate genomes using a custom *S. suis*-specific database (Seemann, 2014). To create this database, ncbi-datasets-cli (v15.24.0) was used to download the .gbff files of all complete genomes of *S. suis* (taxid: 1307). The .gbff files were concatenated and the resulting protein sequences were specified as reference database in prokka using the “--proteins” flag.

### Population structure and phylogenomic analysis

*In silico* multi-locus sequence typing (MLST) was performed using mlst (v2.23.0). Unassigned MLST profiles and new alleles under the “ssuis” scheme were submitted to PubMLST for the assignment of MLST profiles and novel sequence types (ST) (Jolley et al., 2018). *In silico* serotype prediction was carried using the Athey *et al*. pipeline (Athey et al., 2016). PopPUNK (v2.6.4) was used to cluster the *S. suis* genomes, an approach that uses variable-length k-mer comparisons to distinguish closely related isolates and assign them to genomic lineages (Lees et al., 2019). The “poppunk_assign” command was used with the existing *S. suis* database v1 (Murray et al., 2023). Subsequently, “poppunk_visualise” was used to generate a neighbour joining tree, which was visualised in Microreact (Argimón et al., 2016). We next clustered orthologous genes and generated core gene alignments with Panaroo (v1.5.1) using default parameters (Tonkin-Hill et al., 2020). IQ-TREE (v2.4.0) was used to construct a maximum likelihood phylogeny based on the concatenated filtered core gene alignment from Panaroo (Minh et al., 2020). In IQ-Tree, the “-m MFP” parameter was used to identify the best-fit substitution model, “GTR+F+I+R5”, and branch support was assessed with 1,000 ultrafast bootstrap replicates and with “-alrt 1,000”. The resulting tree output was annotated and visualised in Rstudio (v4.4.2) using the R packages ggtree, ggplot2, treeio, ggtreeExtra and ggnewscale. SNP calling was performed using Snippy (v4.6.0) (https://github.com/tseemann/snippy), and recombination regions, defined as loci with elevated densities of base substitutions, were identified and masked using Gubbins (v3.4.0), which constructs a phylogeny based on putative point mutations outside these regions (Croucher et al., 2015).

### Anti-viral defence systems identification

DefenseFinder (v2.0.0) and PADLOC (v2.0.0) were used to predict the presence of anti-viral defence systems in bacterial genomes using .gff and .faa outputs from Prokka as inputs (Payne et al., 2021; Tesson et al., 2024). For DefenseFinder, the “-a” flag was used to also predict anti-defence systems in the genomes and prophage sequences. The results from both tools were consolidated by standardising all system names to lower case to avoid mismatches due to case sensitivity. In this context, “types” refer to broader classification of a defence mechanism (such as “rm”, “cbass”, “crispr-cas”), while “subtypes” indicate distinct variants within a type (such as “rm_type_ii”, “cbass_ii”, and “cas_type_ii”). PADLOC outputs were reformatted to match the structure of DefenseFinder outputs. Defence systems detected by both tools were merged based on overlapping protein components, with subtype conflicts resolved using predefined rules below. Redundant or conflicting entries were deduplicated based on genomic coordinates and protein overlap (Docherty et al., 2025). In cases where the same subtype was labelled with different but equivalent names (such as “cbass_i” vs “cbass_type_i”, and cas_type_i-c vs cas_class1-subtype-i-c), one name was selected arbitrarily to standardise the final annotation. CRISPR-Cas subtypes and arrays in *S. suis* genomes were predicted using CRISPRCasTyper (v1.8.0) (Russel et al., 2020). Detected spacers were dereplicated using CD-Hit (v4.8.1) at identity threshold of 1.0 to obtain unique spacer sequences. Spacer-prophage interactions within and among genomes were examined using SpacePHARER (v5-c2e680a) “predictmatch” module (Zhang et al., 2021). A bipartite network was constructed using the python library “networkx” based on spacer-prophage matches identified with SpacePHARER, which was then visualised in Gephi (v.0.10.1).

Coinfinder (v1.2.1) was used to identify statistically significant co-occurrence and dissociation of defence systems, within the *S. suis* pangenome (Whelan et al., 2020). The newick tree file from IQ-Tree was used as input and the resulting networks of dissociating and associating systems were visualised in Gephi (v.0.10.1), with node size scaled to betweenness centrality.

### Prophage detection, annotation and phylogeny

The “end-to-end” command in geNomad (v1.11.0) was used to detect prophages harboured in the *S. suis* genomes (Camargo et al., 2024). The initial quality assessment of identified prophages was carried out using CheckV (v1.0.3), and only prophages with estimated completeness >50% were included for further analysis (Nayfach et al., 2021). Gene prediction was performed using the Pharokka pipeline (v1.7.5), which integrates PHANOTATE and Prodigal, followed by functional annotation with MMSeqs2 against the PHROGs, VFDB, and CARD databases (Bouras et al., 2023). To improve phage annotations, .gbk files from Pharokka were subsequently analysed with Phold (v0.2.0), a structural homology-based tool that utilises Foldseek and ColabFold (Bouras, 2024). The annotated prophages were further assessed for completeness based on encoded structural genes. A prophage was categorised as full-length if it encoded at least one gene in each of the following modules: “head”, “connector”, and “tail”. Prophages that did not meet this criterion were categorised as incomplete.

Pairwise intergenomic similarity of full-length prophages was computed using VIRIDIC (Moraru et al., 2020). The resulting matrix and cluster tables were used to construct a hierarchical clustering heatmap with ComplexHeatmap in Rstudio (v4.4.2). We attempted to assign the prophages in this study to the lowest possible taxa with taxMyPhage and PhaGCN (v2.1.11) (Millard et al., 2025; Shang et al., 2021). For phylogenomic analysis of prophages, publicly available phage genomes and taxonomy information were downloaded from the INPHARED database (May 2025) (Cook et al., 2021). The nucleotide sequences of all phages that infect bacteria of the *Streptococcus* genus were extracted from the database and combined with the full-length prophages from this study, which were then used as input for the command line version of ViPTreeGen (v1.1.3) with default parameters (Nishimura et al., 2017). The output newick file was annotated with host species; where such information was missing in the database, the host species of were manually searched using the phage accession IDs on GenBank. The tree was visualised in Rstudio using the ggtree, ggplot2, treeio, ggtreeExtra and ggnewscale packages.

### Data wrangling and statistical analysis

Data wrangling was performed using python packages “pandas” and “SeqIO”, and the R package “tidyverse”. Group comparisons were performed using the Wilcox rank-sum test via the ggpubr package, with significance annotated as *p*-values in plots or in figure legends. Summary statistics including median and interquartile range were calculated using dplyr. Correlations were computed using Spearman’s rank correlation coefficient via base R (cor.test, method = “spearman”). Linear models in plots were fitted using the lm() function to evaluate the direction and strength of association. Fisher’s exact test was used to assess the differences in spacer targeting between full-length and incomplete prophages. The results were validated using a chi-squared test for equality of proportions with Yates’ continuity correction (Sahai & Khurshid, 1995). Statistical significance was defined as *p <* 0.05, and this threshold was applied throughout the study.

## Results

### Description of *S. suis* isolated from diseased pigs in Ireland

In total, 112 presumptive *S. suis* isolates recovered from pig swab and post-mortem tissue between 2005 and 2022 were analysed, the majority collected in recent years including 2019 (n=31), 2020 (n=23), and 2021 (n=19) (Table S1). Isolates were initially selected based on alpha-haemolytic activity (24 hours) on sheep’s blood agar and subsequently confirmed as *S. suis* by MALDI-ToF MS prior to sequencing. Following sequencing, two isolates were found to be non-*suis* streptococci and were removed from the dataset. Geographically, isolates originated from 19 counties across Ireland, with the most represented counties being Cork (n=20), Kildare (n=12), Carlow (n=9), and Kilkenny (n=9). Assembly quality statistics for all isolates are provided in Table S1.

### Serotypes and Sequence Types of Irish Isolates

Serotyping of 110 confirmed *S. suis* isolates was carried out using both multiplex PCR and *in silico* approaches. Final serotypes were assigned by combining both methods, prioritising the method with a non-ambiguous result. Fifteen distinct serotypes were identified along with 8 untyped (unknown) isolates. Serotype 9 (n=35) and serotype 2 (n=30) were the most dominant (Fig. 1A). There was generally strong agreement between multiplex PCR and *in silico* typing. However, the PCR-based method could not discriminate between serotypes 1 and 14 or 2 and 1/2. In those cases, *in silico* typing provided resolution, confirming the final designation as serotype 14 or 2, respectively.

**Fig. 1:**
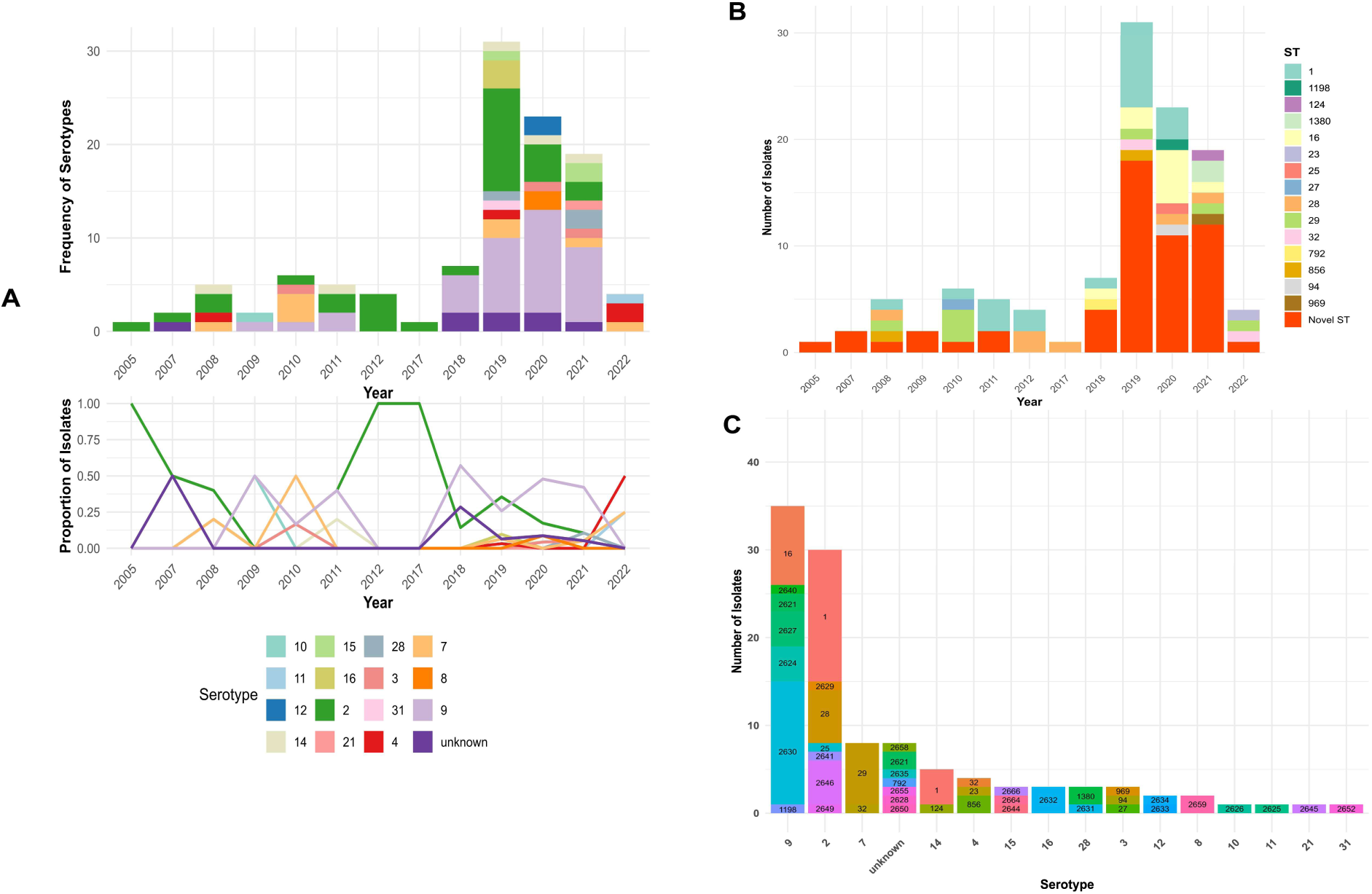
Temporal and genotypic distribution of S. suis isolates by serotype and sequence type. **(A)** Top: Stacked bar plot of frequency of isolates and serotypes between 2005 to 2022. Bottom: Proportional distribution of serotypes over time. **(B)** Distribution of STs over time. All new STs identified in this study are grouped as “Novel ST”. (C) Distribution of STs across serotypes.

*In silico* MLST revealed high genetic diversity among the 110 isolates, with a total of 41 distinct STs identified. Initially, 55 of the isolates could not be matched to any known STs within the existing *S. suis* MLST schemes. Further analysis revealed 18 novel allelic variants of the seven housekeeping genes (*aroA*=2, *cpn60*=1, *dpr*=3, *gki*=2, *mutS*=4, *recA*=5, *thrA*=1). These new allelic profiles were submitted to PubMLST and have been validated, subsequently allowing the assignment of 26 novel STs (ST2621, ST2624-ST2635, ST2640, ST2641, ST2644-ST2646, ST2649, ST2650, ST2652, ST2655, ST2658, ST2659, ST2664, ST2666). ST1 was the most frequently encountered ST (n=26) followed by ST2630 (n=12) while other STs appeared in lower numbers. Novel STs were detected intermittently in early years but became increasingly common from 2018-2021, where they accounted for >50% of cases compared to previously known STs (Fig. 1B). However, this trend could be a reflection of the higher sampling intensity in recent years.

A highly heterogenous distribution of STs was observed across serotypes (Fig. 1C). Serotype 9 was associated with 7 distinct STs, with ST2630 and ST16 accounting for 65.7% of all serotype 9 isolates. Similarly, 8 distinct STs were associated with serotype 2, with ST1 and ST28 accounting for 60% of all serotype 2 isolates. Seven of the eight untypable (serotype) isolates corresponded to novel STs.

Based on clinical presentation and the anatomical origin of samples, the isolates were categorised into pathotypes. Respiratory isolates were recovered from tissue including lung and tonsils and were implicated in infections such as pneumonia and rhinitis. In contrast, the systemic isolates were recovered from neurological or systemic sites such as brain, cerebrospinal fluid, joint, and heart, which were linked to invasive infections such as meningitis, endocarditis, and arthritis. Of the 110 isolates, 54.5% were classified as systemic, 38.2% as respiratory, and 7.3% had insufficient data and were designated as unknown (Table S1).

### Phylogeny and Population Structure of Irish Isolates in the Global *S. suis* Landscape

Given the limited resolution of MLST, which is based on only seven housekeeping genes, PopPUNK was used to capture whole-genome variation and provide a higher resolution view of strain diversity and phylogenetic relationships. This approach facilitated the assignment of all Irish isolates to global lineages and allowed assessment of the isolates within a reference database of 2,424 *S. suis* genomes representing the central population of the species. The central population consists of a genetically cohesive group of lineages with maximum pairwise distance of 0.08 difference per nucleotide in core genome alignment (Murray et al., 2023). The global PopPUNK-based phylogeny illustrates the distribution of local Irish isolates across the central population. Local isolates were not restricted to a small number of clades but were rather distributed across diverse branches within the tree (Fig. 2A). Several local isolates fell into divergent low-frequency lineages, which contributes to the expansion of known population diversity within the central population of *S. suis*.

**Fig. 2:**
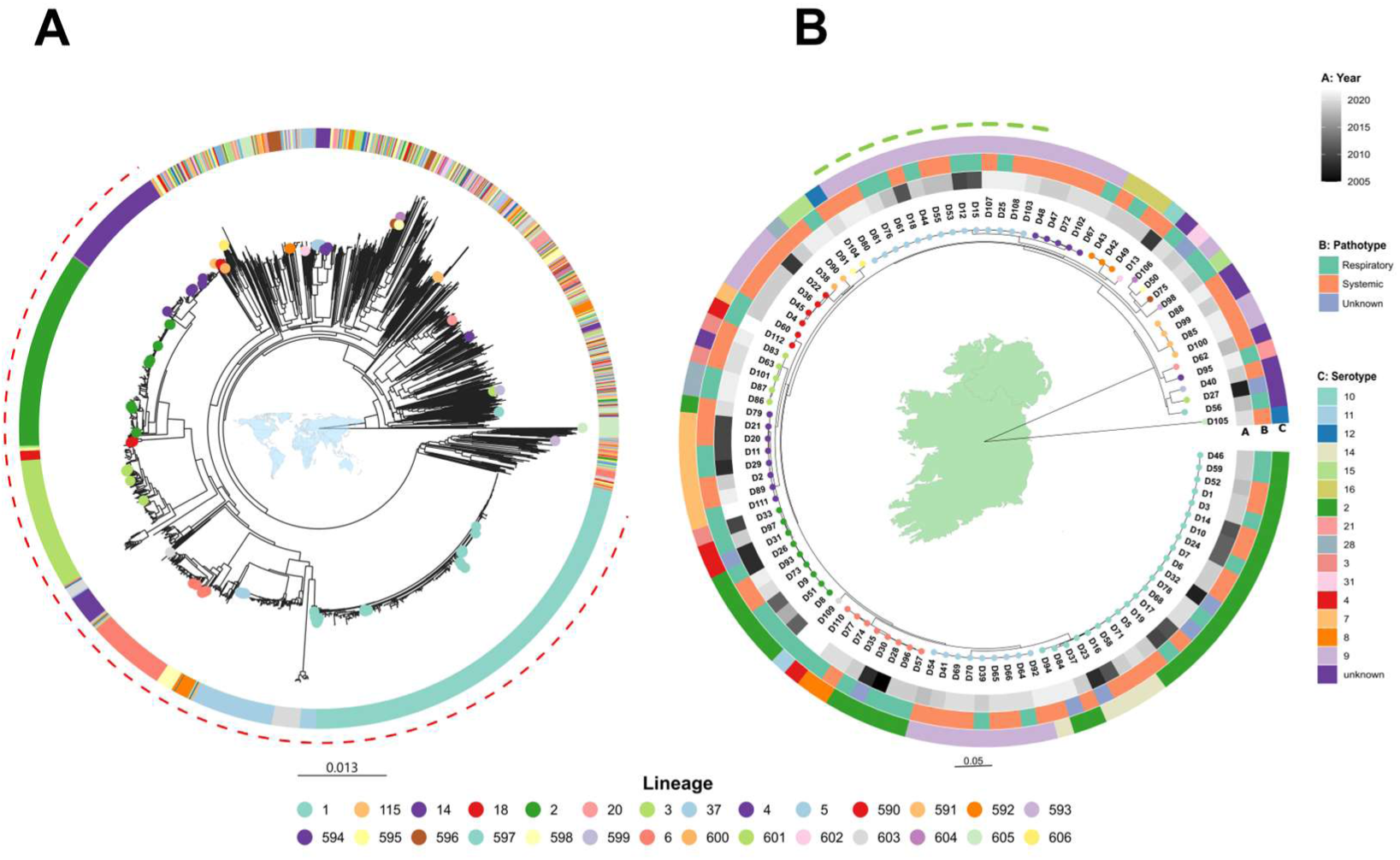
Inferred phylogenetic relationships between S. suis isolates recovered from Irish farms and global sources. **(A)** Phylogenetic placement of Irish isolates recovered from diseased pigs within the global context, based on PopPUNK lineages. A circular neighbour joining tree was constructed using core distances. The tip nodes of only the genomes of Irish origin are coloured. The outer ring represents the assigned lineages for both Irish and global isolates. The dashed red arc indicates isolates within the ten pathogenic lineages of the central population of S. suis. **(B)** A midpoint-rooted maximum likelihood tree inferred for the Irish isolates using IQ-Tree from an alignment of core genes (n=1284) identified by Panaroo. Tip nodes are coloured based on PopPUNK lineage assignment. The dashed green arc represents Irish isolates within lineage 37. The scale of each tree represents nucleotide substitutions per site.

In the national context, Irish isolates were assigned to 28 distinct lineages, which reflects substantial genomic heterogeneity within the collection (Table S1). Among the 28 lineages, 16 were singletons, each represented by only one isolate. Additionally, 25 isolates were assigned to 17 potentially novel lineages not previously defined in the global *S. suis* PopPUNK database. More than half (56.4%) of the Irish isolates clustered within the pathogenic lineages (lineage 1-10) of the central population. Of the Irish isolates within the pathogenic lineages, majority (23 of 62) belonged to lineage 1, which largely corresponds to ST1 isolates (19 of 23). Beyond the pathogenic lineages, lineage 37 was the most common, with 57% of its isolates associated with systemic disease. Overall, several other lineages displayed varying proportions of systemic diseases (Table S2), however, no lineage was significantly enriched for systemic cases after correction for multiple comparisons.

To explore the local context in detail, a maximum-likelihood phylogeny of the isolates was inferred with IQ-Tree based on genome alignments of 1284 core genes generated using Panaroo. The resulting tree revealed a highly resolved, diverse population structure consistent with a polyphyletic population (Fig. 2B). While the population structure was broadly polyphyletic, a notable exception was observed in lineage 37, the most prevalent among the Irish isolates beyond the defined pathogenic lineages (lineages 1-10). We investigated the structure of this lineage by generating a recombination-masked whole-genome SNP phylogeny using Snippy and Gubbins, with the Irish isolate D12 as reference due to its early isolation (2010) compared to the other Irish lineage 37 isolates (Fig. 3A). A total of 19 strains group within lineage 37, of which 14 were from our Irish dataset and belonged to serotype 9 and ST2630, spanning 2010 to 2021 suggesting a sustained local presence. The remaining five strains were isolated in USA and Canada. Most Irish isolates showed no evidence of recombination and harboured very few SNPs (0–36), indicating high sequence conservation (Table S3). One outlier, D18, had 1,251 SNPs, 1,239 of which were located within two recombination blocks (spanning 64,659 bp). However, upon inspection, this block was attributed to the integration of a 65 kb putative conjugative plasmid. Comparatively, three of the five non-Irish lineage 37 genomes had undergone several recombination events (30–78 blocks) resulting in 2,837–6,509 SNPs, whereas one had no detectable recombination and only 73 SNPs, suggesting greater genomic similarity to the Irish cluster. These findings were supported by pairwise average nucleotide identity values, where Irish isolates shared >99.95% ANI (Fig. 3B, Table S4). In comparison, ANI between Irish and non-Irish genomes within lineage 37 ranged from 99.31% to 99.73%, with D18 consistently showing greatest divergence, in line with its recombination associated variation.

**Fig. 3:**
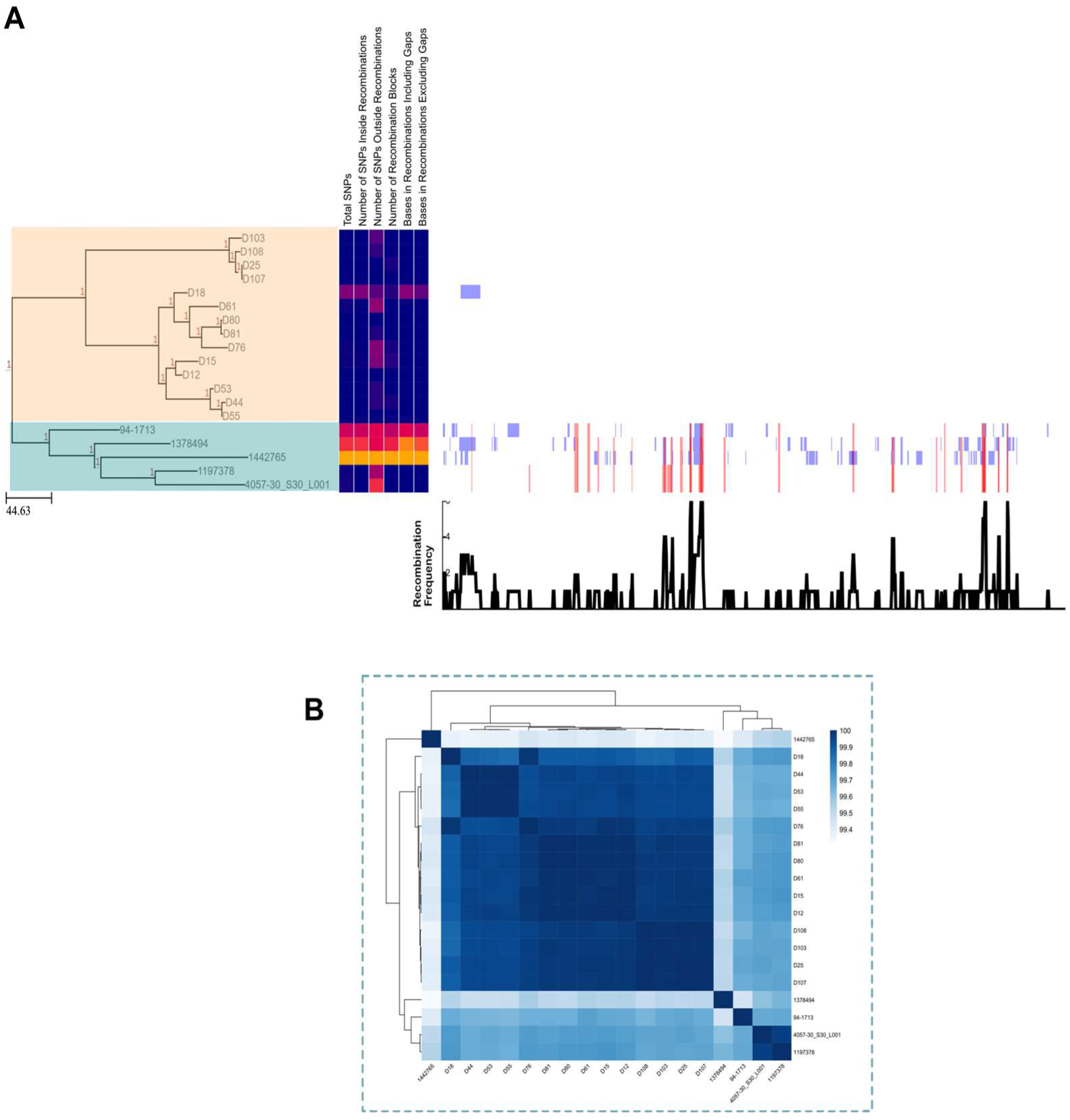
Phylogeny and recombination analysis of S. suis PopPUNK lineage 37. **(A)** Maximum likelihood phylogeny inferred from a recombination-masked whole-genome alignment using Snippy and Gubbins. The tree was visualised in Phandango. The peach-coloured clade represents Irish isolates, while teal denotes previously described non-Irish lineage 37 isolates (Murray et al. 2023). Recombination statistics per branch (Table S3), including total number of SNPs and number of recombination blocks, are presented in a heatmap (blue = low, red = high). In the annotations on the right side, blue represents unique recombination events in individual isolates, while red blocks denote shared/ancestral events. Scale bar represents the number of nucleotide substitutions (SNPs) per site across the recombination-masked alignment. **(B)** Heatmap with dendrogram showing hierarchical clustering of all lineage 37 isolates based on pairwise average nucleotide identity.

### *S. suis* genomes harbour several prophages

Given the central role of mobile genetic elements in shaping bacterial genomes, we examined the incidence and extent of prophage carriage in the genomes of the Irish isolates to explore their contribution to genomic plasticity and divergence. A total of 297 prophage regions with varying degrees of completeness were predicted from 99/110 genomes. The 11 genomes without prophages showed no obvious associations with specific serotypes, STs, or geographic locations. After QC with CheckV and filtering to remove regions of ≤50% completeness, 197 high-confidence prophage regions were obtained across 96 host genomes. Each of these 96 genomes, harboured between 1 to 6 prophages, with an average of 1.7 prophages per bacterial genome (Fig. 4A).

**Fig. 4:**
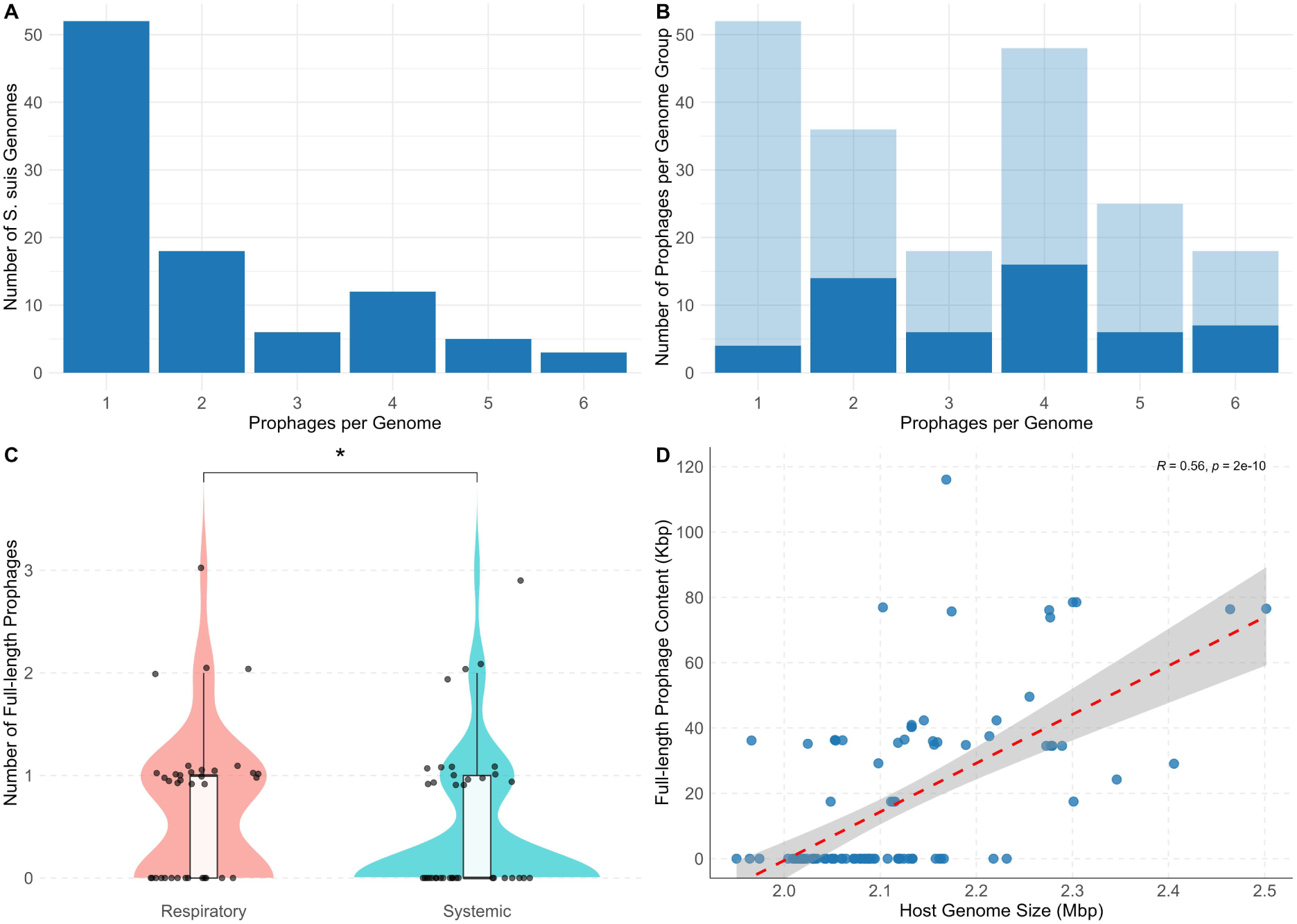
Distribution of prophages in S. suis genomes. **(A)** Prophage counts per genome among isolates that harbour ≥1 prophage (n=96 host genomes). **(B)** Distribution of prophage completeness. Genomes were grouped by the total prophage count (x axis: 1–6 prophages per genome), and all prophages within each group were pooled and counted (y-axis). Bars are stacked by prophage completeness: full-length (dark blue) and incomplete (light blue). The proportion of full-length prophages remain variable across different prophage loads, with no correlation observed between prophage load and the number of full-length prophages (Spearman’s ρ = 0.35, p = 0.49). **(C)** Distribution of full-length prophages between respiratory and systemic isolates. Each point represents a genome, with its vertical position indicating the number of full-length prophages it carries. Points are softly jittered along the y-axis to improve visibility at overlapping values. Respiratory isolates harboured slightly more full-length prophages than systemic isolates (Wilcox p = 0.021). **(D)** Positive correlation between host genome size and total full-length prophage sequence (Kbp) per genome (Spearman’s ρ = 0.0.56, p < 0.001). Dashed red line denotes fitted linear regression. Spearman’s correlation coefficient (R) is displayed.

The prophages were then divided into two categories: full-length or incomplete prophages. Full-length prophages (n=53), defined as prophages that encoded at least one gene in each of the head, connector, and tail structural modules and thus predicted to be complete. The remaining prophages (n=144) were classified as incomplete. Full-length prophages had genome sizes ranging from 17,464 bp to 49,578 bp (median = 35,964 bp) while genomes of incomplete prophages (n=144) ranged from 6,281 bp to 41,118 bp (median = 10,645 bp) in size. When genomes were grouped by the total prophage load (1–6), there was no correlation between total prophages per genome and the number of full-length prophages, suggesting that higher prophage counts do not necessarily reflect a proportional increase in full-length prophages (Spearman’s ρ = 0.35, *p* = 0.49, Fig. 4B). In contrast, a positive correlation was observed between host genome size and total prophage content (Kbp) (Spearman’s ρ = 0.56, *p* < 0.001), indicating larger genomes tend to encode more prophage DNA (Fig. 4D).

To determine whether full-length prophage burden differed between pathotypes, we compared prophage counts per genome in respiratory and systemic isolates. There was a difference between the two groups (Wilcox rank-sum test, *p* = 0.021), with respiratory isolates carrying a slightly higher median number of full-length prophages (median = 1, IQR = 1) compared to systemic isolates (median = 0, IQR = 1) (Fig. 4C). Nonetheless, considerable overlap in distribution was observed between the two groups, indicating that although the difference is statistically significant, it may be modest in biological terms.

### Diverse *S. suis* phages are embedded across the phylogeny of streptococcal phages

To explore the sequence diversity and relatedness of the full-length prophages, VIRIDIC was used to estimate intergenomic similarities. Prophages were grouped into genus and species clusters using a nucleotide identity threshold of 70% and 95%, respectively, in accordance with the ICTV standard for dsDNA viruses. In total, 28 genera and 34 species clusters were identified within the 53 full-length prophages with pairwise similarities ranging from 0% to 100% (Fig. 5, Table S5). This clustering delineated multiple groups of closely related prophages as well as numerous distinct prophages. For instance, in species cluster 9, which comprises eight members, six of them shared >99.9% identity. Despite being identified in different host genomes, these prophages are essentially identical, suggesting the circulation of a single prophage across multiple host backgrounds spanning different years (2005–2022), counties, serotypes (2 and 4), and STs (ST23 and ST2646). Conversely, many prophages were genomically unique. The genus-level clustering identified 28 distinct clusters, however, of these clusters, 16 were singletons. Of the remaining clusters, eight consisted of two members, two clusters possessed four members, 1 cluster possessed five members and one cluster had eight members. Attempts to assign the prophages to established taxa using TaxMyPhage and PhaGCN were unsuccessful below the class level, which suggests that all 53 full-length prophages fall outside currently defined phage taxonomic groups (below class level). Thus, the prophages likely belong to 28 novel genera, spread across 34 novel species.

**Fig. 5:**
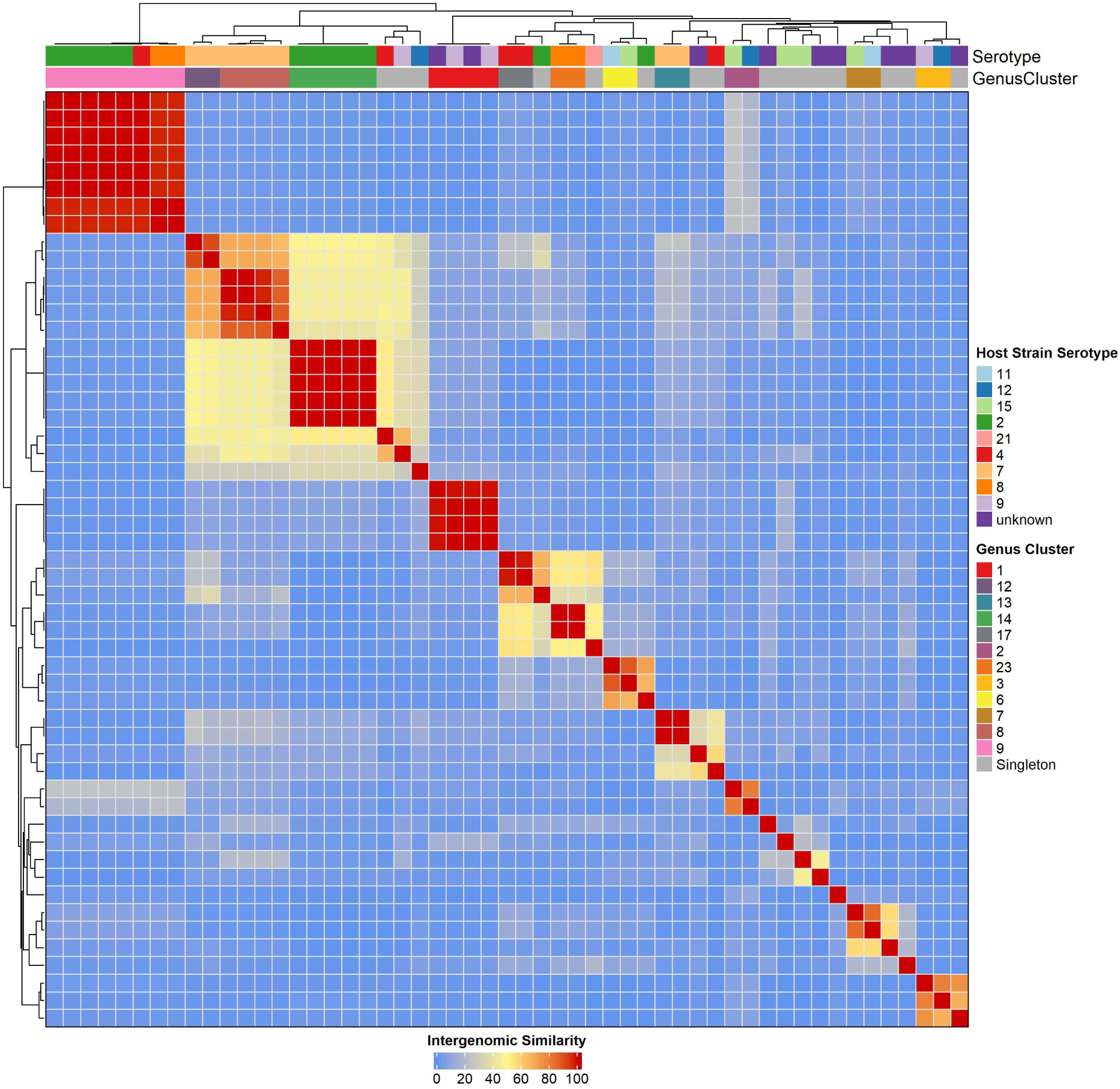
Intergenomic similarity of full-length prophages. Full-length prophages were classified into genus clusters according to ICTV genus delineation of 75% shared nucleotide identity.

To place the full-length prophage genomes of *S. suis* into perspective, a viral proteomic tree was constructed based on tBLASTx computations of genome-wide similarities of all known phages (n=780) that infect streptococcal species (n=49) together with the full-length prophages from this study (Table S6). Of the 780 publicly available streptococcal phage sequences, 83 were associated with *S. suis*, largely from uncharacterised deposited prophage sequences (Rezaei Javan et al., 2019). Only four of the publicly available *S. suis* prophages were placed in an established sub-family, Trabyvirinae, suggesting that most *S. suis* phages either belong to novel viral taxa or represent members of highly divergent, poorly sampled taxa. In general, the *S. suis* phages (including those from this study), do not cluster randomly in the phylogeny but rather form multiple distinct clades, most of which are composed exclusively of *S. suis*-specific phages (Fig. 6). The full-length prophage sequences from this study were interspersed with those of publicly available *S. suis* phages, indicating that they expand and reinforce previously under-sampled clade rather than defining entirely novel branches of the viral tree. A similar trend was observed for phages of other pathogenic streptococci such as *Streptococcus pyogenes* and *Streptococcus agalactiae*, which demonstrated broader phylogenetic dispersion and formed several, often poorly resolved clades with limited taxonomic assignments. Contrastingly, phages from non-pathogenic streptococci—in particular, *Streptococcus thermophilus*, a dairy-associated commensal— formed tightly clustered, monophyletic clades almost exclusively composed of *S. thermophilus*-infecting phages. Additionally, these phages have been extensively characterised taxonomically into well-defined genera including *Moineauvirus*, *Brussowvirus*, *Piorkowskivirus* and *Vansinderenvirus* (Fig. 6). Although most *S. thermophilus* phages are phylogenetically distinct from those infecting pathogenic species including *S. suis*, *S. pyogenes*, and *S. pneumoniae*, phages belonging to the genus *Vansinderenvirus* form a clade that is more closely related to several *S. suis* phages than to other *S. thermophilus* phages, albeit involving distant relationships overall.

**Fig. 6:**
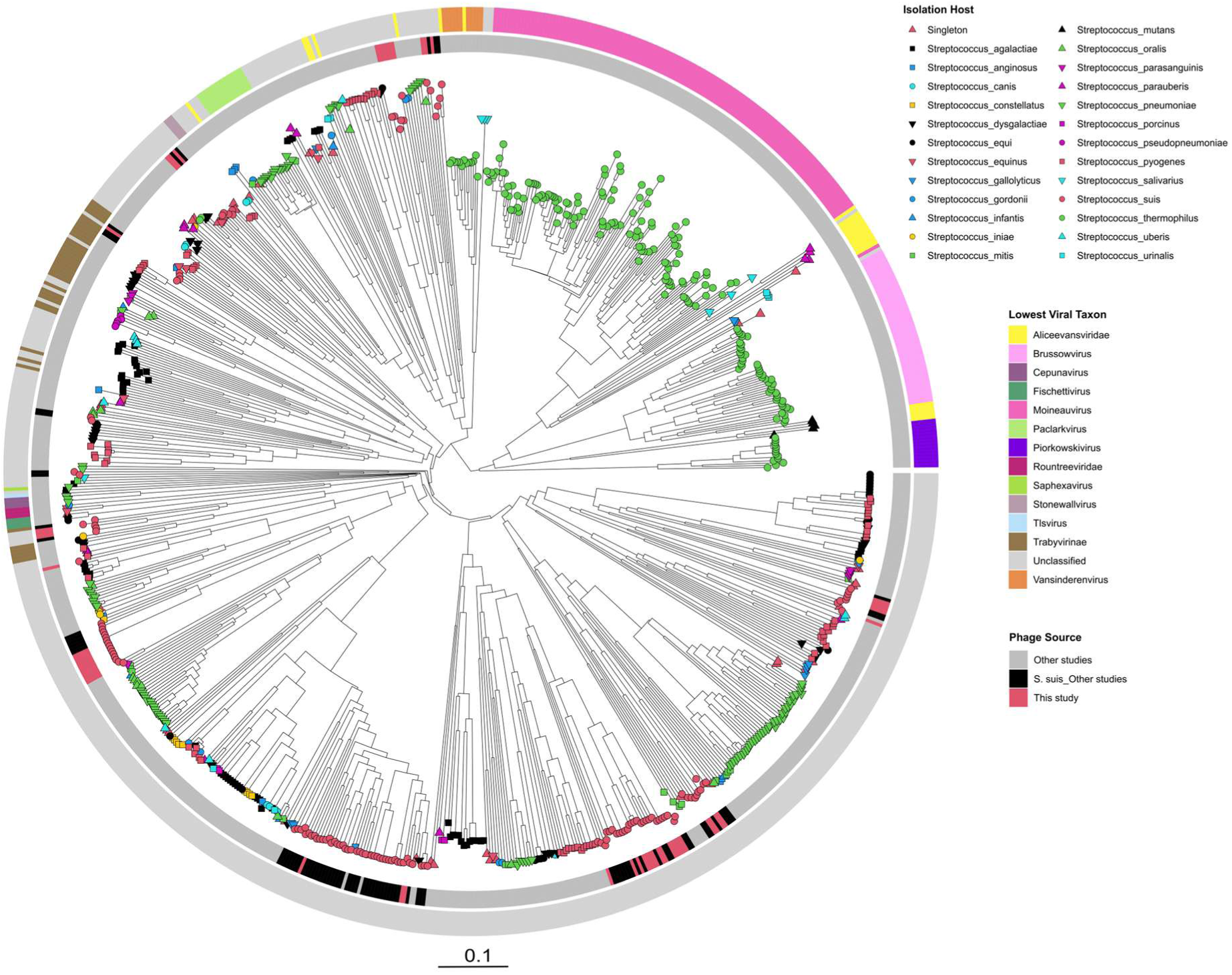
Viral proteomic tree of streptococcal phages, including full-length prophages identified in this study. Tip nodes are annotated by shape and colour to indicate the species of the host bacterium, with S. suis shown in red circle, S. thermophilus in green circle, and other streptococcal species in additional shapes and colours. The inner ring indicates source of phages, including S. suis prophages identified in this study (red), S. suis prophages from other studies (black), and all other non-S. suis streptococcal phages from other studies (grey). The outer ring represents the lowest viral taxonomic classification, including the genus Vansinderenvirus (orange). Although most S. thermophilus phages (green circle tip node) from distinct clades, members of Vansinderenvirus cluster more closely with several S. suis phages (red circle tip nodes) than with other S. thermophilus phages, despite the relationship being relatively distant overall.

### Defence systems are prevalent in *S. suis* genomes

After deduplication and consolidating the predictions from PADLOC and DefenseFinder, a total of 1,394 defence systems were identified in the *S. suis* genomes. These predicted hits could be classified into 69 distinct subtypes (Table S7). Here, broader defence mechanism categories including “rm”, “cbass”, “crispr-cas” are designated as “types”, while specific variants within each category such as “rm_type_ii”, “cbass_ii”, “cas_type_ii” are classified as “subtypes”. Each of the 110 genomes harboured between 5 to 25 system occurrences, with an average of 12.67 ± 3.63 systems (mean ± SD). In terms of diversity, 5 to 17 unique subtypes were identified per genome with a median of 11 (Fig. 7A). On average, respiratory isolates encoded more defence systems (median = 13.5) than systemic isolates (median = 11) (*p* = 0.016). Similarly, a difference was observed in the density of defence systems, with respiratory isolates encoding 6.33 systems/Mbp of host genome compared to 5.35 defence systems/Mbp in systemic isolates (*p* = 0.028, Fig. 7B). A strong positive correlation was observed between bacterial genome size and the number of defence systems they encode (Spearman’s ρ = 0.796, *p* < 0.001, Fig. 7C), indicating that isolates with larger genome sizes are likely to encode more defence systems. A similar trend was identified, with a positive correlation between the number of prophages and the defence systems encoded per genome (Fig. 7D, Spearman’s ρ = 0.6, *p* < 0.001).

**Fig. 7:**
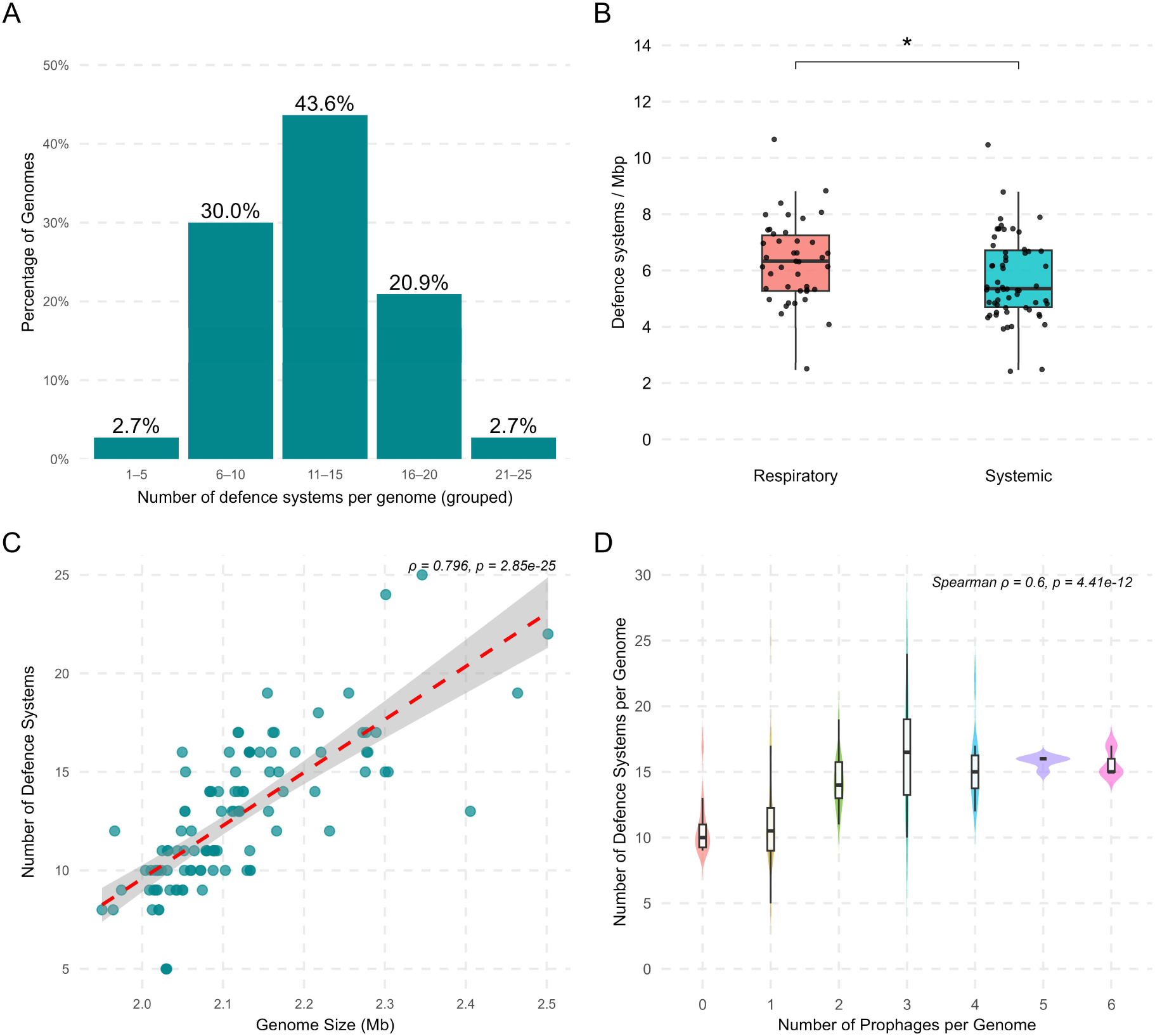
Prevalence of defence systems in S. suis. **(A)** Proportion of genomes that encode a number of unique defence systems. Genomes encoded 5 to 17 (median = 11) unique defence subtypes. **(B)** Density of defence systems per Mbp of host genome. Density of defence systems was significantly higher in respiratory isolates (median = 6.33) compared to systemic isolates (median = 5.35) (Wilcox p = 0.028). **(C)** Correlation between host genome size and number of defence systems. A strong positive correlation was observed between (Spearman’s ρ = 0.796, p < 0.001). **(D)** Association between number of prophages per genome and abundance of defence systems.

### Diverse defence systems are abundant and often co-occur in *S. suis* genomes

The 69 distinct defence system subtypes were classified into 29 broader types, including all PDC (PADLOC defence candidate) subtypes under PDC and all PD (“Phage Defense” hits identified by DefenseFinder) subtypes grouped under “Other” as they represent unvalidated systems or those with unknown mechanism of action (Table S7). All isolates encoded at least one PDC-type system, while PD-type systems were present in 99.1% of isolates. Among the validated systems, restriction modification (RM) systems were the most prevalent system type, present in 97.3% of genomes, followed by abortive infection (Abi) systems (74.6%), RosmerTA (55.4%), and DMS (47.3%). In contrast, CRISPR-Cas systems were identified in only 26.4% of isolates belonging to different genetic backgrounds with no clear association to specific lineages, STs, or serotypes (Fig. 8A).

**Fig. 8:**
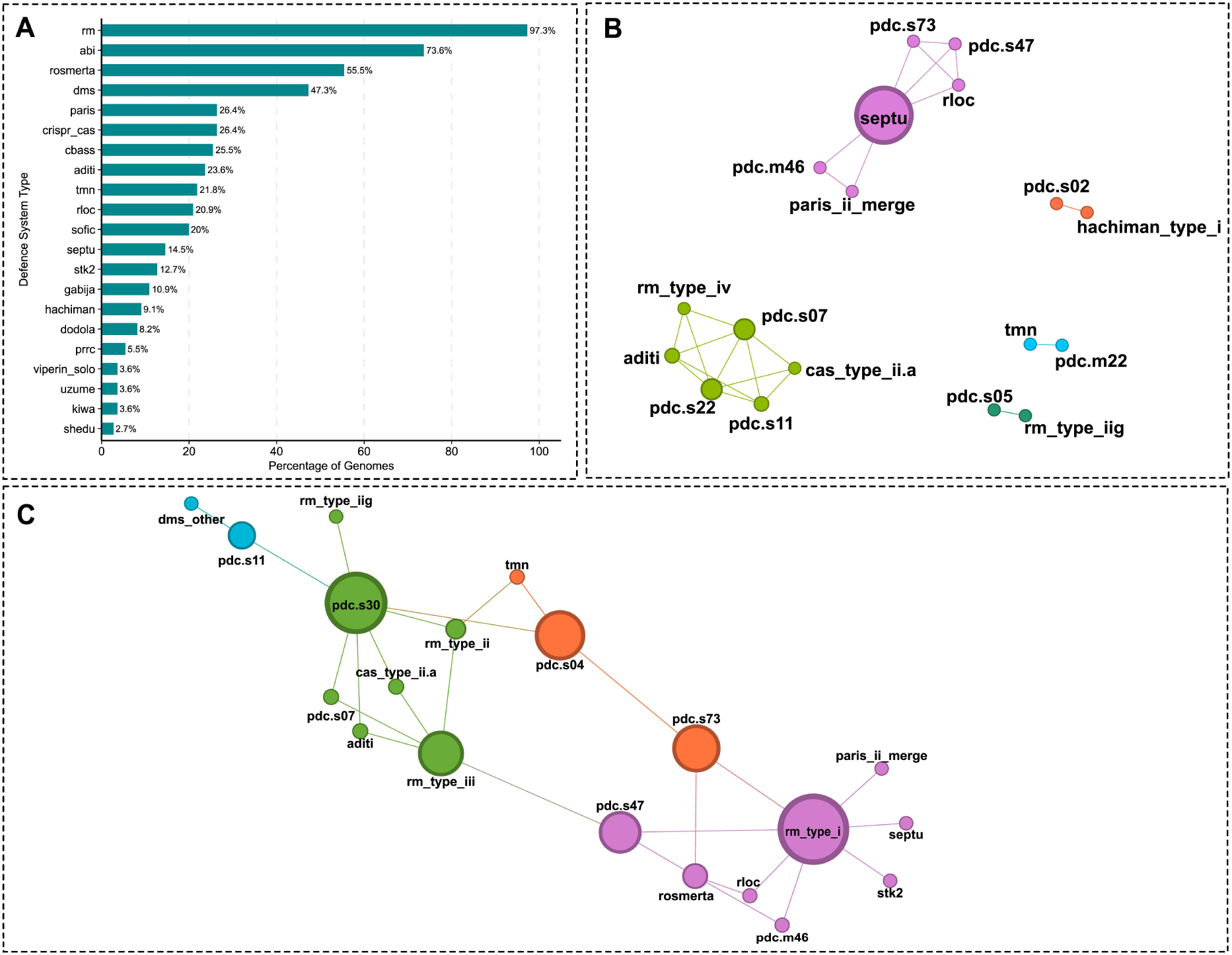
Prevalence and co-occurrence patterns of defence systems in S. suis genomes. **(A)** Prevalence of defence systems across all genomes. Bars represent the percentage of genomes encoding each validated defence type ( excludes those in the “Padloc Defence Candidates” and “Phage Defense” categories). Network of statistically co-occurring **(B)** and dissociated **(C)** subtypes identified using Coinfinder (includes the unvalidated PDC and PD systems). Nodes represent individual subtypes; edges indicate co-occurrence or dissociating events that were more frequent than expected by chance. Node size in panels B and C is scaled to betweenness centrality. Node colours in B and C represent communities identified by modularity-based community detection in Gephi.

At the subtype level, RM type I was the most abundant with a total of 156 occurrences (median per genome = 1.71) across 82.7% of isolates. PD-T4-6, although slightly less abundant (109 occurrences), was the most prevalent defence subtype, present in 99.1% of all isolates (Table S7). Other abundant subtypes include PDC-S04 and RM Type II, which were detected in 55.4% (116 occurrences) and 62.7% (79 occurrences) of isolates, respectively. In contrast, 31 of the 69 subtypes including Cas Type I-C, AbiG, AbiK, ietAS, Thoeris, and DRT Class II were detected in <5% of genomes. Among the 1,394 defence systems identified across all genomes, only 10 were found to be encoded by prophages, including both full-length and incomplete prophages. These systems, representing six unique subtypes (AbiD, AbiZ, Hachiman, Lamassu-family, RM Type II, and RM Type III), were identified within nine distinct prophages.

To assess organisational patterns of the detected subtypes across the genomes, we used Coinfinder to identify statistically significant co-occurrences and mutual exclusion events that occur frequently or less frequently than what would be expected by chance within the pangenome. A total of 24 subtype pairs were found to positively co-occur while 27 subtype pairs were detected as dissociated (*p* < 0.05, Bonferroni-corrected, Table S8). Numerous PDC-subtypes displayed strong associations, such as PDC-S07 with PDC-S11, PDC-S22, and RM Type IV (Fig. 8B). Similarly, the recently discovered Aditi system was associated with PDC-S07, PDC-S11, PDC-S22, and RM Type IV. In addition, CRISPR-Cas Type II-A also positively co-occurred with PDC-S07, PDC-S11, and PDC-S22. Among the 27 pairs of dissociating subtypes, PDC-S07, PDC-S11, and Aditi were found to dissociate with PDC-S30, as well as RM Type I with both stk2 and Septu (Fig 8C). To investigate whether co-occurring defence subtypes were also physically clustered within genomes, we estimated genomic distance based on whether co-occurring pairs were also within 20 genes of each other. Of the 24 pairs, only one, PDC-S07 and RM Type IV were consistently co-localised (23/23 occurrences).

We investigated whether anti-defence systems (ADS), which are employed by prophages to counter host defences, were encoded in the genomes of the isolates. A total of 167 ADS were identified across the 110 genomes, with an average of 1.7 ± 0.88 per genome (median = 1). Ninety-eight of the 110 genomes encoded at least one ADS. These systems could be grouped into two distinct subtypes: the anti-CRISPR subtype acriia21, which was detected in 87.3% of genomes and accounted for 153 of the 167 total ADS identified; and the anti-RM subtype ardaardu, found in 12.7% of genomes (n=14). Also, only 5 of the 167 ADS were found to be encoded in prophage regions.

### CRISPR-Cas adaptive immunity and phage interactions in *S. suis*

Although CRISPR-Cas defence systems were detected in only 26.4% of genomes, anti-CRISPR ADS were identified in 87.3% of the genomes. This imbalance suggests that despite the limited prevalence of CRISPR-Cas systems, they may exert enough selective pressure to drive the widespread maintenance of anti-CRISPR systems. To examine this further, we used CRISPRCasTyper and SpacePHARER to investigate the architecture and diversity of CRISPR arrays with focus on spacer-prophage interactions. In agreement with predictions from PADLOC and DefenseFinder, CRISPRCasTyper identified 37 complete CRISPR loci in 26.4% (n=29) of the genomes which belonged to two subtypes, CRISPR-Cas II-A (n=26) and CRISPR-Cas I-C (n=3). A total of 363 spacers were identified in 34 genomes, ranging from 2 to 33 spacers per array. Clustering the spacers at 100% sequence identity yielded 182 unique spacers, indicating that nearly 50% were shared within or between genomes. To explore the tripartite relationships among genomes, spacers, and prophages, we identified 198 spacer-prophages targeting events involving 41 unique spacers found in the genomes of 12 isolates (Fig. 9A). These spacers targeted 43 out of the 197 prophages identified in this study. Targeting by spacers was strongly biased toward full-length prophages (75.5% of 53) compared to only 3 out of 144 (2.1%) incomplete prophages. This enrichment was highly statistically significant (Fisher’s exact test, *p <* 0.001; odds ratio = 136.7, Fig. 9B) and was confirmed by a two-sample test for equality of proportions (*p <* 0.001; 95% CI: 60.3–86.5%). We explored spacer-phage relationships further using the 780 publicly available streptococcal (pro)phages against the spacers identified in Irish isolates in this study. Of the 363 spacers and 780 phages, a total of 755 spacer-phage events were identified involving 278 unique phages and 41 spacers. These phages targeted by the spacers infect 32 of the 49 streptococcal species represented in the streptococcal phage database, including *S. suis*. Interestingly, the majority of the targeting events were directed against phages that infect *S. thermophilus* (n=126), followed by *S. suis* (n=33), *S. pneumoniae* (n=22), and *S. pyogenes* (n=20). Additionally, all three of the previously isolated phages of *S. suis* including Bonnie, Clyde, and SMP were among the targeted phages with 11, 2, and 8 matching spacers, respectively.

**Fig. 9:**
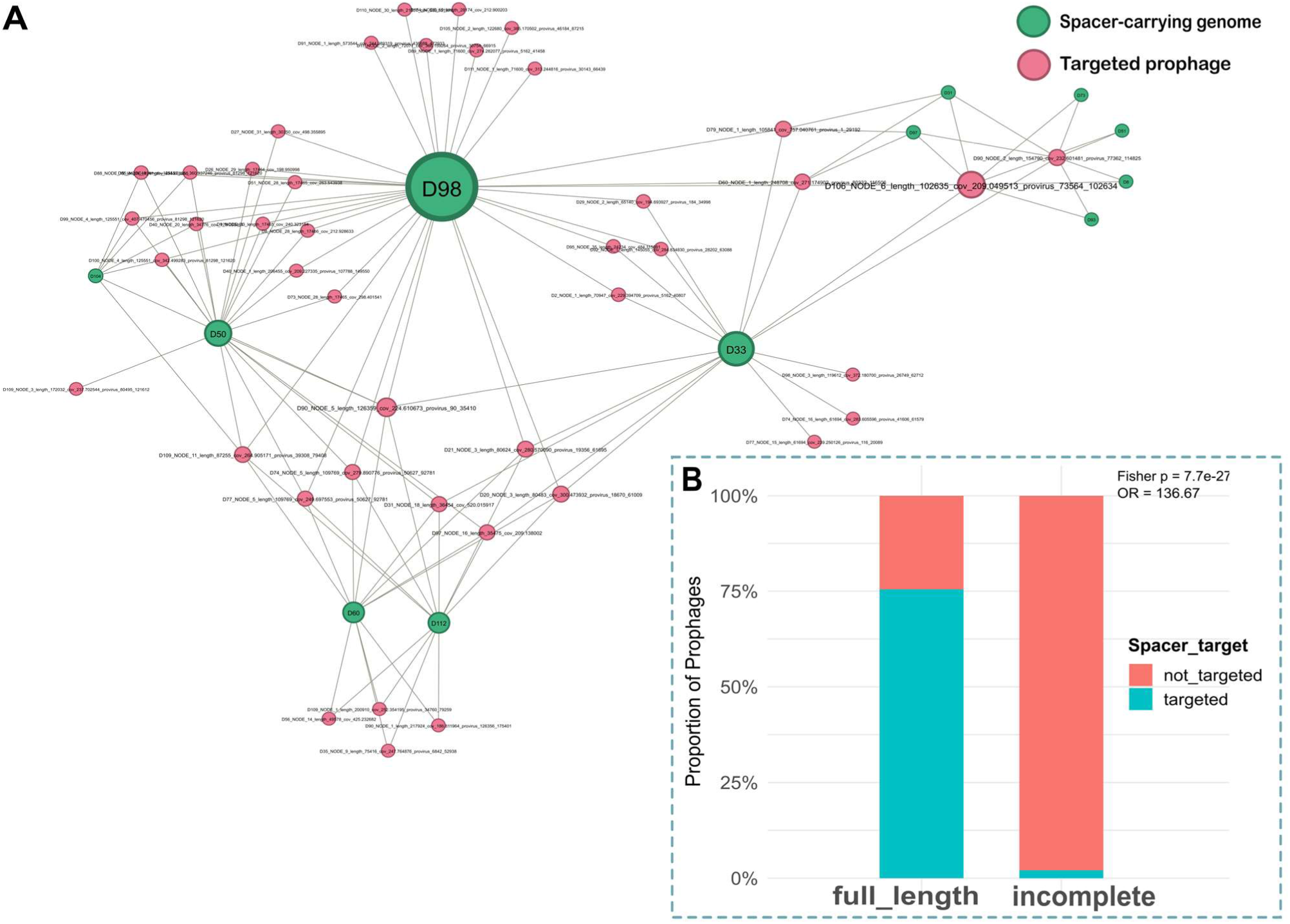
CRISPR spacer-prophage interactions in S. suis genomes. **(A)** Network of spacer-prophage interactions. A total of 198 spacer-prophage interactions were detected across 12 genomes, involving 41 unique spacers that target 43 prophages. Nodes in green represent Irish S. suis genomes that encode CRISPR spacers, while pink nodes represent prophages that are targeted by at least one of the spacers. Edges represent spacer-prophage matches identified using SpacePHARER. The network was visualised in Gephi. **(B)** Proportion of full-length and incomplete prophages that were targeted by spacers. Spacer targeting was strongly biased toward full-length prophages (75.5%) versus incomplete prophages (2.1%) (Fisher’s exact test, p < 0.001; odds ratio = 136.7).

## Discussion

In this study, we present the first comprehensive genomic investigation of *S. suis* in the Republic of Ireland. The genomic characterisation of the 110 *S. suis* isolates from diseased pigs collected between 2005 to 2022 reveals a diverse national-level population embedded within the broader global context. Serotypes 9 and 2 were identified as the most dominant serotypes within the isolate collection. This finding is in line with trends reported across western European countries, where serotype 2 was historically the most common cause of disease in pigs, however, over the past two decades, serotype 9 has risen markedly in prevalence and is emerging as the dominant serotype (Goyette-Desjardins et al., 2014). Beyond the two leading serotypes, we identified 14 other serotypes at lower frequencies, including serotypes 7, 14, 4, and untypable isolates. The serotype heterogeneity identified herein mirrors the diversity reported in recent European surveillances (Li et al., 2024). MLST of the Irish isolates showed an equally diverse population including several genotypes related to known worldwide clonal complexes. In addition, 26 previously unreported STs including ST2630, were detected in our relatively small sample set. The discovery of these novel STs in Ireland possibly reflects the high underlying genomic diversity of *S. suis* and a historical sampling bias whereby strains from several pig-producing countries, including Ireland, have been underrepresented in global MLST database compared with their pig industry output, particularly on a per capita level. An open population structure has been described for *S. suis*, characterised by a large accessory genome and frequent horizontal gene transfer leading to continual emergence of new allelic profiles as seen among our collection (Estrada et al., 2022). From a practical perspective, this means that while traditional MLST is useful for first-line categorisation, it may struggle to keep pace with *S. suis* diversity (>2,800 STs). The difference between a novel and a described ST may be a point mutation or gene recombination, yet such differences could have a significant epidemiological relevance. For instance, in our dataset, even within common serotypes like 2 and 9, multiple STs were identified. Both serotyping and MLST offer limited resolution (Lees et al., 2019). Serotyping captures only capsular polysaccharide types and can be confounded by capsular switching through recombination, while MLST targets just seven genes (<1% of the genome). Consequently, isolates may be grouped together if they share similar allelic profiles even if they differ significantly in accessory genome or virulence. Similarly, these methods may separate clonal strains into different STs or serotypes if a single gene mutates or is associated with a recombination event.

To overcome these limitations, we placed the Irish genomes into a global phylogenomic context using PopPUNK and a database of 2,424 *S. suis* genomes (Fig. 2B). This WGS approach showed that the Irish isolates are phylogenetically dispersed throughout the broader global *S. suis* population, rather than clustering in a single clonal group. Murray *et al*. delineated the global population structure of *S. suis*, showing a central population which includes ten PopPUNK pathogenic lineages (1–10). Consistent with that model, more than half of the Irish isolates fell within the ten pathogenic lineages (Murray et al., 2023). For instance, we identified lineage 1 isolates (n=23), which is a highly virulent clade associated with systemic infections and most human cases globally (Dong et al., 2021). The presence of lineage 1 isolates on Irish farms is epidemiologically significant as it confirms this predicted zoonotic clade circulates in Irish pigs. This poses a potential occupational risk to farmers and abattoir workers. While reports of zoonotic infection are more common in Asia, human cases are becoming increasingly reported in Europe. In Ireland, the first documented case occurred in 1995 and involved fatal sepsis in a swine farmer (Baddeley, 1995). Additional cases have been reported in 2014 and 2015 (Brizuela et al., 2024).

Beyond the ten pathogenic lineages, lineage 37 was the most prevalent in the Irish dataset. Further analysis of this clade using recombination-masked SNP phylogeny revealed a tight cluster of 14 Irish isolates belonging to serotype 9 and ST2630 (Fig. 3A). These isolates were collected from geographically distant farms over more than a decade and showed low SNP variation (0–36) and minimal recombination, indicating a stable endemic clonal lineage within Ireland. In contrast, the 5 non-Irish lineage 37 genomes harboured substantially more SNPs and recombination events, indicating a more distant relationship and potentially representing the non-clonal pool from which the Irish lineage 37 isolates could have descended. Pairwise ANI supported this structure, with Irish lineage 37 isolates sharing >99.95% identity, while showing more distant similarity of 99.31–99.73% to non-Irish isolates. Furthermore, while associations between lineage and disease presentation were explored, no lineage was statistically enriched for systemic or respiratory disease after correction for multiple testing. Overall, the presence of the stable endemic lineage alongside other global pathogenic lineages suggests ongoing evolutionary processes that could facilitate the emergence of new variants with enhanced virulence or antimicrobial resistance. These findings highlight potential challenges for disease management on Irish pig farms and necessitates continued genomic surveillance to mitigate risks to both animal health and food safety.

To further contextualise the evolutionary dynamics of the *S. suis* population, we investigated the relationship between prophages and anti-viral defence mechanisms. Prophages that mediate horizontal gene transfer and influence virulence, together with defence systems that shape susceptibility to phages, modulate genome plasticity and evolutionary trajectories. These interactions could also impact the abundance of *S. suis* within microbial communities and ecological success. Understanding these dynamics is not only crucial to pathogen evolution but also has important implications for developing phage-based therapeutic interventions against zoonotic pathogens.

Our analysis revealed nearly all genomes encoded at least one prophage, with several identified as polylysogens, consistent with our previous observations using publicly available *S. suis* genomes from 11 countries (Osei et al., 2022). In comparison, the prophage load is higher than has been reported in other streptococci, including *S. thermophilus*, *S. mutans* and *S. pneumoniae* (Rezaei Javan et al., 2019). We identified a mix of full-length prophages with complete structural modules as well as incomplete degraded prophage remnants in the genome. Although respiratory isolates carried slightly more full-length prophages on average than systemic ones (Fig. 4C), there was considerable overlap in distribution between the groups, which indicates prophage carriage is widespread across pathotypes. Clustering of prophages based on intergenomic similarity revealed 28 and 34 genus and species clusters, respectively (Fig. 5). None of the prophages could be assigned to any known viral taxa below the class level and thus represent members of novel genera and species. This observation is not surprising, given that until recently, only one “lytic” phage had been characterised and sequenced for *S. suis* (Ma & Lu, 2008; Osei et al., 2025). When placed in the context of all known streptococcal phages, the prophages identified in this study interspersed among the described phages rather than forming a single clade (Fig. 6). This finding highlights potential gene flow and shared ancestry with phages of other streptococcal hosts and is consistent with observations that cross-species transmission of prophages occurs among streptococci (Rezaei Javan et al., 2019). Indeed, two of the *S. suis* phages clustered closely with phages infecting *S. anginosus* and *S. orisratti*, which indicates that these viruses or their modules move between host species.

In parallel, we profiled the anti-viral defences encoded by the *S. suis* isolates. Despite their small genomes, *S. suis* encode an average of 12.67 defence systems, which is significantly more than the 5–7 systems found in bacteria with larger genomes like *E. coli* (∼5Mbp), but comparable to the high defence load (12.13) in *Bifidobacterium* species (Docherty et al., 2025). RM systems were nearly ubiquitous in the Irish *S. suis* genomes (97.3%), which is similar to prevalence levels previously reported in *S. suis* (90.2%) and *S. thermophilus* (100%, n=27) but higher than the average for prokaryotes (83%) (Kelleher et al., 2024; Osei et al., 2022; Tesson et al., 2022). Although traditionally viewed as anti-viral defence, Type I RM systems are increasingly recognised as multifunctional, contributing to gene regulation, MGE stabilisation, and virulence under epigenetic control (Manso et al., 2014; Willemse & Schultsz, 2016). Phase-variable Type I and III RM methyltransferases segregate by clade, with lineage-restricted Type I system confined to the zoonotic *S. suis* Bayesian population (BAPS) group 1 and likely acquired at its origin (Atack et al., 2018). These systems may therefore influence both defence variation and the emergence of host-adapted lineages.

The diversity of the defence arsenal extended across several validated systems, such as Abi, RosmerTA, DMS. However, the majority of the detected systems were present in only a small minority of the genomes (Fig. 8A). Interestingly, 24 instances of positively co-occurring pairs were identified including numerous PDCs and RM Type IV, suggesting possible co-selection or synergistic activity (Tesson & Bernheim, 2023; Wu et al., 2024). However, only one of those pairs was co-localised. Conversely, some pairs were mutually exclusive in the same genome, perhaps due to functional redundancy or fitness cost when combined (Mayo-Muñoz et al., 2023; Tesfazgi Mebrhatu et al., 2011). Further experimental validation is needed to elucidate these patterns.

Similar to the trends observed for prophage carriage, respiratory isolates encoded a higher density of defences systems than systemic isolates. In general, there was a positive correlation between the number of defence systems in a genome and both its size and prophage content. Thus, as *S. suis* genomes accumulate MGEs including phages, they also accrue defence systems. This parallel accumulation raises interesting questions about the long-term bacteria-phage arms race. A possible explanation is that larger genomes provide more “genomic real estate” to accommodate prophages and defence systems. A study from 2016 demonstrated that prophage frequency increases with genome size, particularly in genomes <6Mbp, where insertional burden is less constrained by essential gene density (Touchon et al., 2016). This may partly explain why respiratory isolates, which tend to have larger genomes, have higher defence and prophage content. Another factor could be the selective benefits conferred by temperate phages through carriage of accessory functions, including virulence factors, auxiliary metabolic genes, and immunity to superinfecting phages (Owen et al., 2021). Moreover, many prophages themselves encode defence mechanisms including toxin-antitoxin, BstA abortive infection system, and RM systems, which enhances host immunity (Brenes & Laub, 2025; Owen et al., 2021). However, in our dataset, only 10 out of the 1,394 defence system occurrences were associated with prophage regions, highlighting that in *S. suis*, defence systems are largely maintained independently of prophage integration. Finally, the co-occurrence and occasional co-localisation of defence systems may contribute to the accumulation of multiple, complementary systems.

We conducted a detailed analysis of CRISPR-Cas loci and anti-CRISPR systems in the genomes. We observed an interesting paradox wherein, despite only 26.4% of genomes encoding a CRISPR-Cas system, 87.3% carried at least one anti-CRISPR (Acr). However, only 5 of 153 anti-CRISPR were prophage-encoded, suggesting that majority are largely maintained by non-prophage elements. Acr genes could have been acquired horizontally via plasmids or integrative conjugative elements that use Acr proteins to overcome CRISPR defences and facilitate transfer between hosts (Marino et al., 2018; Pinilla-Redondo et al., 2020). These observations suggest that Acr systems may contribute to the evolutionary dynamics between mobile genetic elements and host defence systems beyond prophage-mediated interactions. The relatively low prevalence of CRISPR-Cas systems (subtypes II-A and I-C) is consistent with reports in *S. suis* but significantly lower than *S. thermophilus* or *S. mutans*, in which the system is nearly universally encoded (Kelleher et al., 2024; Osei et al., 2022; Shields et al., 2020). Notwithstanding, several spacers were identified among strains that encode a CRISPR system. There was a substantial spacer targeting bias towards full-length, presumably inducible prophages rather than incomplete remnants (Fig. 9). This suggests that these full-length prophages are infective and the CRISPR spacers are retained by the cell to prevent successful infections. It also implies that phage infection is a constant threat for *S. suis*, despite limited successes in isolating infective phages in a lab setting. For example, we found that several spacers in *S. suis* strains target the isolated *S. suis* phages SMP, Bonnie, and Clyde (Ma & Lu, 2008; Osei et al., 2025). The presence of spacers homologous to regions in these phages provides evidence that the Irish strains or their ancestors encountered these phages or closely related phages. In fact, we previously reported that Bonnie and Clyde infect some of these strains, thus it is noteworthy that some strains carry spacers against them, likely a result of historical phage infection events (Osei et al., 2025). More broadly, numerous spacers shared sequence identity with non-*S. suis* streptococcal phages including those that infect *S. thermophilus*, *S. pneumoniae*, and *S. pyogenes*. While this may suggest cross-species exposure or shared phage pools, it is unlikely due to host-specific phage receptor recognition and the low intergenomic similarity between *S. suis* phages and those of other streptococci (Kortright et al., 2020). However, the phylogenetic clustering of phages belonging to the genus *Vansinderenvirus* with *S. suis* phages rather than other *S. thermophilus* phages, despite overall distant relationships, provides compelling evidence of evolutionary connections between these phage lineages. This relationship, combined with early observations that *Vansinderenvirus* representative, 5093, possesses a mosaic genome, lacks canonical anti-repressor genes, and shares high homology with prophages of non-dairy streptococci, suggests that these spacer matches capture genuine evolutionary remnants (Mills et al., 2011). The poorly sampled and highly divergent nature of *S. suis* phage taxa may represent missing evolutionary links that help explain these cross-species phylogenetic relationships.

## Conclusion

This study provides the first comprehensive genomic characterisation of *S. suis* in Ireland, revealing a diverse population shaped by endemic and global lineages with significant implications for pig production and occupational risk to those working with pigs and handling pork products. The presence of zoonotic clades and novel STs highlights the need for genomic surveillance to track transmission along the farm-to-fork continuum. Our integrated analysis of prophages and defence systems reveals a dynamic mobilome and active phage-host arms race that shapes genome plasticity and may influence ecological success. Furthermore, the full-length prophages offer a promising platform for phage-based biocontrol. The epidemiological and genomic data presented herein offers a robust foundation for future surveillance and intervention efforts within the One Health context.

## Data availability

Raw sequence reads have been deposited in the NCBI Sequence Read Archive and are available under BioProject PRJNA1295417 (https://www.ncbi.nlm.nih.gov/bioproject/PRJNA1295417/)

## Supplementary materials

Table S1 to S7 are available on Zenodo at https://doi.org/10.5281/zenodo.16987951

## Ethics statement

This study was approved by the Teagasc Animal Ethics Committee (Reference: TAEC2022-335). No live animals were experimentally infected for this study.

## Acknowledgements

This project (Improved Pig Health through the Novel Application of SynBio in Phage Therapy, 2020US-IRL201) was funded by the Irish Department of Agriculture, Food and the Marine, through the 2020 US-IRL R&D Partnership Call.

## CRediT authorship contribution statement

**Emmanuel Kuffour Osei:** conceptualisation, data curation, formal analysis, investigation, methodology, visualisation, writing – original draft, writing – review and editing. **A. Kate O’Mahony:** conceptualisation, formal analysis, methodology, writing – review and editing. **Reuben O’Hea:** data curation and investigation. **John Moriarty:** data curation and investigation, writing – review and editing. **Áine O’Doherty:** data curation and investigation. **Margaret Wilson:** data curation and investigation. **Edgar Garcia Manzanilla:** funding acquisition, methodology, writing – review and editing. **Jennifer Mahony:** conceptualisation, funding acquisition and project management, investigation, methodology, writing – original draft, writing – review and editing. **John G Kenny:** conceptualisation, funding acquisition and project management, investigation, methodology, writing – original draft, writing – review and editing.

